# Alterations in neurovascular coupling following acute traumatic brain injury

**DOI:** 10.1101/183129

**Authors:** Hyounguk Jang, Stanley Huang, Daniel X. Hammer, Lin Wang, Meijun Ye, Cristin G. Welle, Jonathan A. N. Fisher

## Abstract

Traumatic brain injury (TBI) is a leading cause of mortality and disability worldwide. A challenge for diagnosing and assessing the severity of TBI, however, is that quantitative biomarkers are lacking. We explored potential functional indicators for TBI by noninvasively monitoring sensory-evoked electrical and hemodynamic activity using a novel hybrid optical and electrophysiological measurement approach. By combining diffuse correlation spectroscopy with co-localized electrophysiological measurements in a mouse model of TBI, we observed concomitant alterations in somatosensory-evoked cerebral blood flow and electrical potentials following controlled cortical impact. Injury acutely reduced the amplitude of stimulus-evoked responses, which mostly recovered to baseline values within 30 min; intertrial variability for these parameters was also acutely altered. The kinetics of recovery, however, varied among specific components of the evoked waveforms, and we observed strong correlations between the two measurement modalities for only a select subset of waveform parameters. Overall, our results identify a novel set of potential biomarkers for TBI and demonstrate the utility of combined, noninvasive optical and electrophysiological measurements for detecting injury-induced abnormalities in neurovascular reactivity.

## Introduction

In the U.S. each year there are over 1.5 million TBIs, resulting in 50,000 deaths [1]. TBI induces temporary or permanent impairment of cognition, physical function, and psychosocial behavior [2]. TBI severity ranges widely depending on the nature of the injury. The majority of these injuries are classified as “mild” TBI (mTBI), which are challenging to diagnose and track because quantitative biomarkers for mTBI are lacking. In fact, one of the defining characteristics of mTBI is that it cannot be validated through standard methods of clinical imaging [3]. Given the difficulty of rapid detection, mTBI poses a particular challenge to public health because repeated injuries such as concussions have a cumulative effect on brain health [4]–[6].

Among a wide spectrum of TBI sequelae, sensory and cognitive deficits are some of the most common [7]. A difficulty in assessing these deficits is that they are frequently mutually confounding. For example, auditory dysfunction following injury includes both peripheral deficits, e.g. increased hearing thresholds, as well as difficulties with more sophisticated auditory tasks, such as discriminating sounds in noisy environments or comprehending speech, (Gallun et al., 2012b). Visual deficits following TBI are similarly complex; the most common complaints, for example, are accommodative deficiencies [8] which include blurred vision, headache, motion sickness, or loss of concentration during visual task performance [9]. While auditory and visual performance may be difficult to dissociate from cognitive deficits, olfaction, which does not involve as significant feedback with subcortical structures, is also impacted by TBI, and the degree of anosmia is correlated with severity of injury [10].

The integrity of sensory systems can be probed electrophysiologically via evoked potentials (EPs). EPs can be measured noninvasively and comprise a series of positive and negative voltage deflections, which reflect the afferent relay and processing of sensory information. Auditory, visual and somatosensory evoked potentials (AEPs, VEPs, and SSEPs, respectively) are routinely used in multiple clinical contexts to aid neurological assessment of TBI and to monitor functional recovery over time (Irimia et al., 2013). Sensory-evoked neural activity can also be inferred from the cerebral hemodynamic response function (HRF), which depicts changes in cerebral blood flow (ΔCBF), volume, and oxygenation that are evoked by brief sensory stimuli. Damage to any portion of the cerebral vasculature fundamentally alters the ability of the network to supply neurons with energy [11]. In fact, blockage of even single capillaries can cause larger-scale changes in blood flow [12] and may result in microvascular ischemia [13]. Repercussions of these injuries, even if mild, may continue to progress following the primary trauma, ultimately leading to more global sequelae.

We explored the possibility of deriving novel indicators for mTBI based on intrinsic correlations between hemodynamic and neuronal activity. To noninvasively monitor sensory-evoked hemodynamics, we used diffuse correlation spectroscopy (DCS), which takes advantage of the dynamic scattering properties of red blood cells to directly measure CBF. DCS is particularly sensitive to flow in the cortical microvasculature due to the high absorption (and thus low probability of photon escape) in larger blood vessels [14]. DCS has been used to measure functional hemodynamics associated with sensory stimuli and motor tasks [15], [16], and has also been used to track baseline CBF following brain trauma [17]. We supplemented our ongoing optical recordings of the sensory-evoked HRF with concomitant measurements of SSEPs and applied this multimodal approach to a mouse model of TBI that employed controlled cortical impact (CCI) as the source of primary injury [18]–[20]. The ability to noninvasively monitor both aspects of neural response enabled us to obtain the first detailed, *in vivo* portrait of the effects of acute injury on sensory processing in the brain.

## Materials and Methods

### Surgical Procedures

All animal experiments were performed in accordance with the guidelines of the White Oak Institutional Animal Care and Use Committee. Optical and electrophysiological measurements were performed on 6 male C57BL/6J mice (12 – 24 weeks). Anesthesia was induced by an initial exposure to 4% isoflurane for less than 30s. Animals were additionally administered an injection of xylazine (18 mg/kg, IP) to provide a stable plane of anesthesia at low isoflurane concentrations (0.1-0.25%) for the remainder of the experimental session, which typically lasted 3-6 hours. Maintenance doses of xylazine (6 mg/kg) were administered once every ~2.5 hours. Following initial anesthesia induction, animals were positioned in a stereotaxic apparatus (David Kopf Instruments, CA). Their body temperature was measured and maintained at 37°C with a closed-loop temperature controlled heating pad (Model TC-1000, CWE). Respiratory rate was also monitored and maintained at ~100 breaths per minute during the surgical and experimental procedures. Skin incisions were infused with lidocaine and the eyes were covered with ointment (Lacri-Lube) to prevent drying. A midline sagittal incision was made in the skin, which exposed the coronal and lambdoid sutures on the skull. The intersection of these sutures with the midline (i.e. bregma and lambda) served as landmarks for recording locations and were also used as a guide when drilling burr holes. The area of the skull under the probe was cleaned with 70% ethanol and the optical probe was secured to the skull by cyanoacrylate glue (Loctite 454, Hankel, Australia) (Fig. 1a). Burr holes (diameter: ~0.5 mm) were made for placement of two silver wire electrodes: a recording electrode, placed 2.5 mm lateral and 1 mm posterior to bregma, and a reference electrode, placed 1 mm lateral and 1 mm posterior to lambda. The recording electrode was embedded within the optical probe tip, and the reference electrode was secured into position with Kwik-Sil adhesive (World Precision Instruments, FL). A ground needle electrode was placed subcutaneously on the back of the animal.

**Fig. 1.**
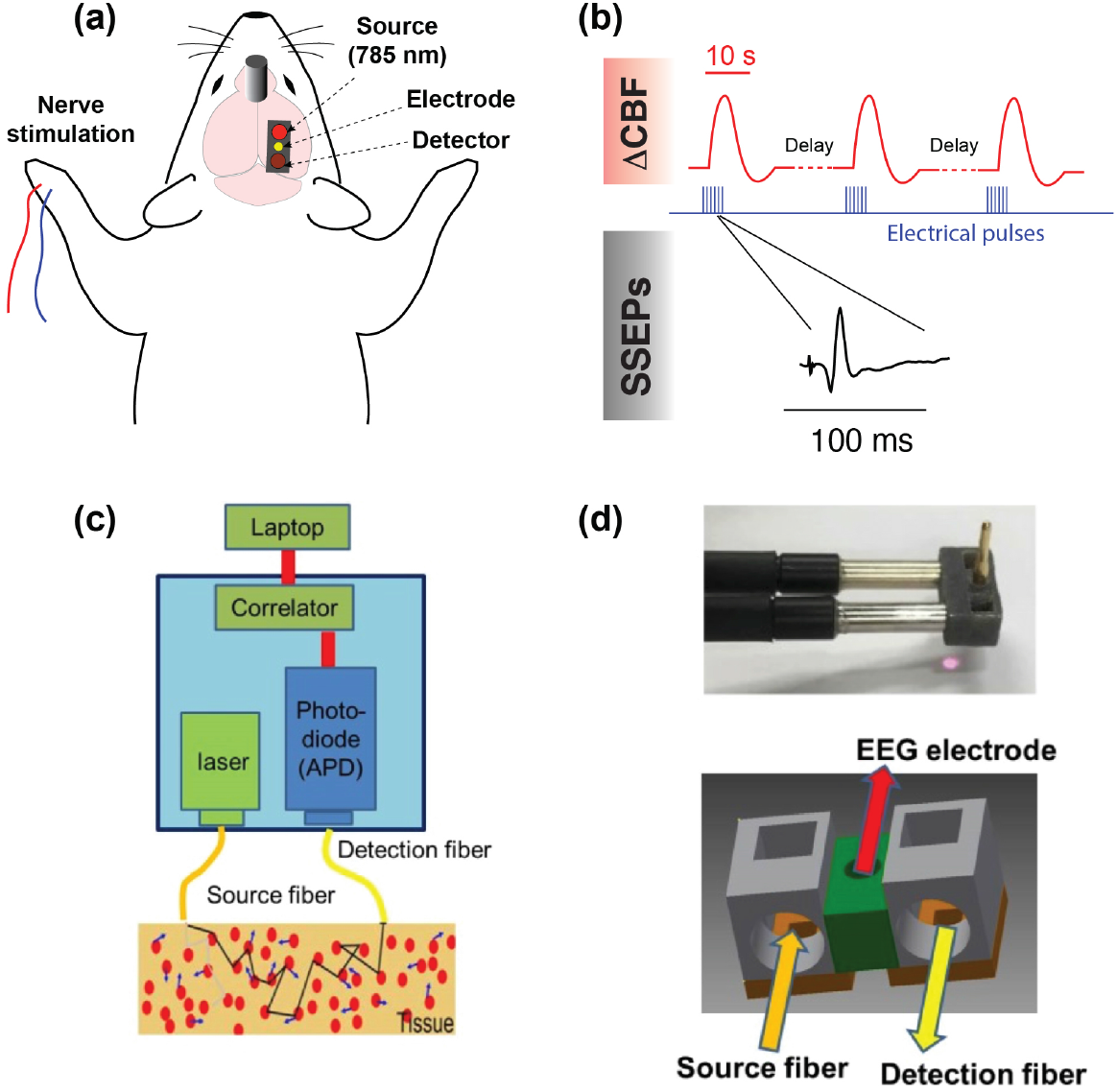
Experimental design and measurement apparatus. (a) Schematic of the recording and stimulation configuration. A DCS probe tip containing optical fibers is placed on the region of the skull above the primary somatosensory cortex, at a location coinciding with the forepaw’s representation. A recording EEG electrode is inserted through the center of the probe. Pulses of electrical current are delivered to the contralateral limb with respect to the optical and electrophysiological measurements. Closed-skull impact is delivered by a 5-mm impactor tip ~4 mm anterior and ~1 mm lateral of bregma contralateral to the optical measurement (see MATERIALS AND METHODS). (b) Overview of synchronization between stimulus, measurements of CBF, and SSEPs. The red traces depict simulated hemodynamic signals, which are broad compared with electrophysiological potentials (note the time scale difference in the insets). The blue pulses indicate electrical stimuli delivered to the median nerve. (c) Block diagram of the DCS device used in this study. (d) Photograph and schematic of the 3D-printed DCS probe tip. Light is delivered and collected from the tissue through fiber optics that are coupled to the skin with micro-prisms.

### Sensory Stimuli and Electrophysiological Recordings

SSEPs were recorded in single-ended configuration (RZ5D processor, PZ2 preamplifier, ZC16 headstage, Tucker-Davis Technologies, FL) with a shared common reference electrode, at a sampling frequency of 3 kHz. Optical measurements of ΔCBF were performed concurrently, driven by a separate computer. The median nerve contralateral to the recording locations was stimulated via a pair of 27-gauge stainless-steel needles inserted subcutaneously into the forelimb of the animal. The electrical stimulus consisted of a train of 12 current pulses generated by a constant current stimulator (DS7A, Digitimer) (amplitude 4 mA, frequency 3 Hz, pulse duration 200 μs, total pulse-train duration 4 s). Individual measurements of sensory-evoked ΔCBF were separated by 45 s, which we empirically found to be the minimum inter-trial duration that did not elicit alterations in the steady-state blood flow index. An overview of the acquisition and stimulus timing is shown in Fig. 1b.

### Controlled Cortical Impact (CCI)

Controlled cortical impact (CCI) is a well-established and highly reproducible brain injury model [18], [19], [21]–[24]. Closed-skull impact was delivered with an Impact One™ stereotaxic impactor (Leica Microsystems). Using a 5-mm diameter metal impact tip, we used the following settings: velocity 4 m/s, dwell time 100 ms, impact depth 0.6 mm. Strike velocities >5 m/s and/or strike depths > 0.8 mm resulted in skull fracture. Additionally, impactor tips with smaller diameter tended to cause fractures. The metal impactor tip was positioned over the exposed skull ~4 mm anterior and ~1 mm lateral of bregma, contralateral to the optical measurement.

### Optical Measurement of Cerebral Blood Flow

The DCS signal can be used to measure blood flow by using speckle correlation techniques. The basic theory underlying DCS has been extensively described in previous publications [14], [25]. Briefly, when laser light migrates through tissue, the emerging intensity pattern, called a speckle pattern, is composed of bright and dark spots, which are caused by constructive and destructive interference of photons that traverse different path lengths. The intensity fluctuations of a single region, which are caused by interactions of scattered light with moving particles (i.e. red blood cells), can be utilized to extract information about blood flow.

A block diagram of our DCS recording system is shown in Fig. 1c. It consists of a long coherence length continuous-wave near-infrared (NIR) laser (785 nm, CrystaLaser, NV), a photon-counting avalanche photodiode (APD) (SPCM-AQRH-12-FC, Excelitas, Quebec, Canada), and an autocorrelator signal processing board (www.correlator.com, NJ). NIR excitation light was delivered to the brain with a multi-mode optical fiber (200 μm core diameter, Thorlabs, NJ) and the scattered light was detected with a single-mode fiber (5 μm core diameter) connected to the APD. The source and detection fibers were separated by 5 mm. The APD signal was sent to the correlator board, which computed the intensity of the autocorrelation function. The correlation board streams output signals continuously via USB to a laptop PC for further data analysis.

The probe that coupled the optical fibers and recording electrode to the head was fabricated from semi-flexible acrylate polymer using a three-dimensional printer (Objet 260 Connex 3 printer, Stratasys, MN). Embedded microprisms directed light from the fibers down to the head. The recording electrode was situated at the midpoint between the DCS optical source-detector separation. The midpoint of the probe (indicated in green in Fig 1d) was thinner and permitted the probe to bend and conform to the curvature of the mouse’s skull, providing tight contact to the surface. The probe was positioned on a relatively flat region of intact skull above the forelimb’s representation in primary somatosensory cortex, 2.5 mm lateral of the midline at the same coordinate as bregma on the rostrocaudal axis (Paxinos 2004).

*Modeling the Hemodynamic Response Function*

To extract quantitative features from the sensory evoked HRF, we fit the observed waveform to a canonical HRF that uses two gamma functions to approximate the hemodynamic response (Fig. 2a). We used a linear combination of two gamma functions [26], i.e.

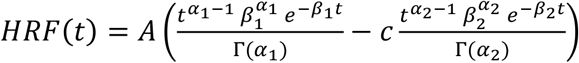

where Г represents the gamma function. A, c, α_1_, α_2_, β_1_, and β_2_ are fitting coefficients, which were fit using the Levenberg–Marquardt algorithm in MATLAB. Here, A controls the amplitude, c determines the ratio of the response to undershoot, and α_1_, α_2_, β_1_, and β_2_ control the shape of the fit function. We obtained four major features, which are the amplitude and temporal delay (relative to stimulus onset) of the HRF’s initial peak and undershoot minimum, to quantitatively describe ΔCBF based on the fitted curve.

**Fig. 2.**
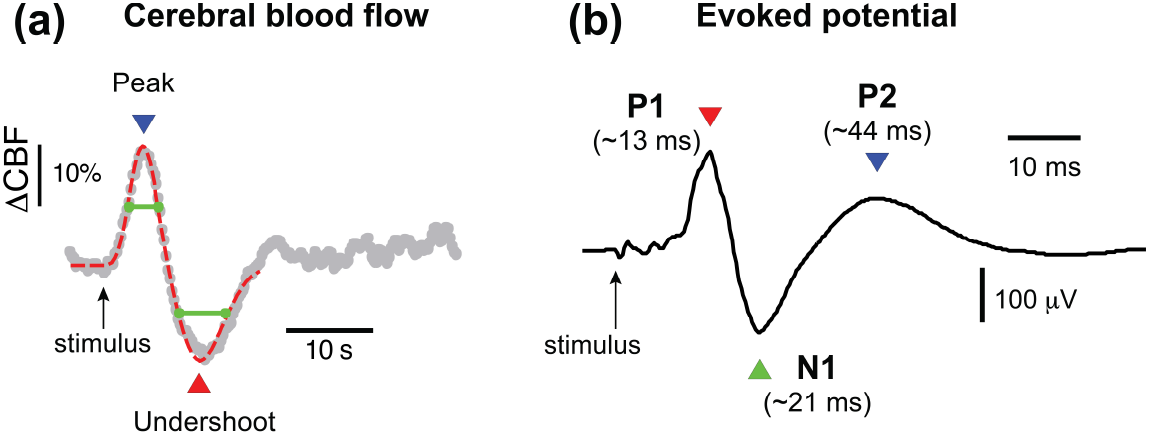
Somatosensory-evoked hemodynamic and electrophysiological responses. (a) Optically-measured changes in relative cerebral blood flow (CBF) following electrical stimulation of the median nerve. The gray trace represents the average of 10 trials in one experiment; blood flow values are normalized to the average of CBF values recorded in the four seconds prior to the stimulus onset, which is indicated by the arrow. The dotted red line that is superimposed on the trace represents a fit to the observed hemodynamic response using a conventional hemodynamic response function model (two gamma functions), as described in MATERIALS AND METHODS. (b) SSEP recorded concurrently during acquisition of the optical data depicted in (a). The prominent peaks P1, N1, and P2 occur at temporal latencies that are well-established in the literature. The trace represents the average obtained from the same 10 trials in (a), however for each trial there are 12 SSEPs recorded in response to individual stimulus pulses (4 s of pulses delivered at 3 Hz), thus the trace represents the average of 120 SSEPs. The green lines represent the widths of positive and negative peaks, measured at half of the maximum / minimum values. Note that the timescale in (a) is roughly 10× longer than in (b).

### Time-Frequency Analysis of the Hemodynamic Response Function

In the interest of parsing apparent oscillatory behavior associated with post-injury modifications in the hemodynamic response following the main positive and negative peaks, we performed spectrotemporal analysis on the CBF data throughout the entire duration of six experiments. Briefly, the continuous DCS data was divided into six five-minute epochs, and analysis of the evoked responses within those epochs was used to inform single “frames.” We utilized time-frequency transform functions from the MATLAB toolbox EEGLAB [27] to produce spectrograms that quantified power as well as inter-trial coherence (ITC), which indicates the degree of phase-locking in the observed HRF. Note that for each data point in Figs. 6 and 7, the spectrograms were based on only the trials within the five-minute epochs; although a larger sample size generally improves signal-to-noise, in this case because the HRF was dynamically changing in time following injury, there was a tradeoff between signal-to-noise and the temporal resolution with which the time evolution of power and ITC could be quantified. We empirically converged on five-minute epochs as a compromise based on observations from the continuous CBF that indicated that recovery generally occurred on a timescale slower than five minutes following a rapid decrement due to CCI.

## Results

### Concurrent Measurements of Sensory-Evoked Cerebral Hemodynamics and SSEPs

Using the DCS component of our multimodal apparatus, we recorded sensory-evoked changes in the HRF concurrently with electrophysiological measurements of SSEPs. Fig. 2a depicts an example of the pre-injury hemodynamic response. Averaged over six animals, the HRF displayed an initial increase in blood flow of 14 ± 4.8% (mean ± SD) that peaked at 9 ± 0.3 s, followed by a subsequent undershoot of magnitude -12 ± 2.5%, which reached a negative peak 17 ± 1.1 s after the stimulus (12 ± 1.1s after the first peak). CBF returned to baseline values roughly 10 s following maximum undershoot. In terms of electrophysiological recordings, SSEPs displayed well-defined peaks, most prominently a positive deflection (P1) at ~12 ms post-stimulus, a negative peak (N1) at ~21 ms, and broader positive peak at ~44 ms (P2) (Fig. 2b, n = 6 mice).

### TBI Acutely Alters Hemodynamic and Electrophysiological Functional Responses

Cortical impact acutely reduced the amplitude and altered the waveforms of sensory-evoked hemodynamic and electrophysiological responses (Fig. 3). Within the first 5 minutes following impact, the peak of the HRF decreased by ~50%, and the undershoot amplitude decreased as well, though to a lesser extent (~20 – 40%). The peak latencies additionally displayed changes on the order of ~10%, though with opposite trends: the initial increase in CBF peaked ~1 s earlier than pre-injury, and the undershoot peaked ~1 s *later* than in the pre-injury case. The alterations in these parameters are summarized in Fig. 4. The peak-to-peak amplitude recovered to approximately 80% of its baseline value within 30 minutes, and during this period, the shifts in peak latencies resolved completely. It should be noted that despite being more sensitive to injury than other features of the HRF, the initial peak also recovered at a relatively fast rate, and within 5 – 10 minutes following injury, the peak amplitude had essentially recovered completely. In contrast, CBF undershoot changes following injury, though more difficult to quantify because of the variability in that portion of the waveform post CCI, displayed no recovery trend within the same time period.

**Fig. 3.**
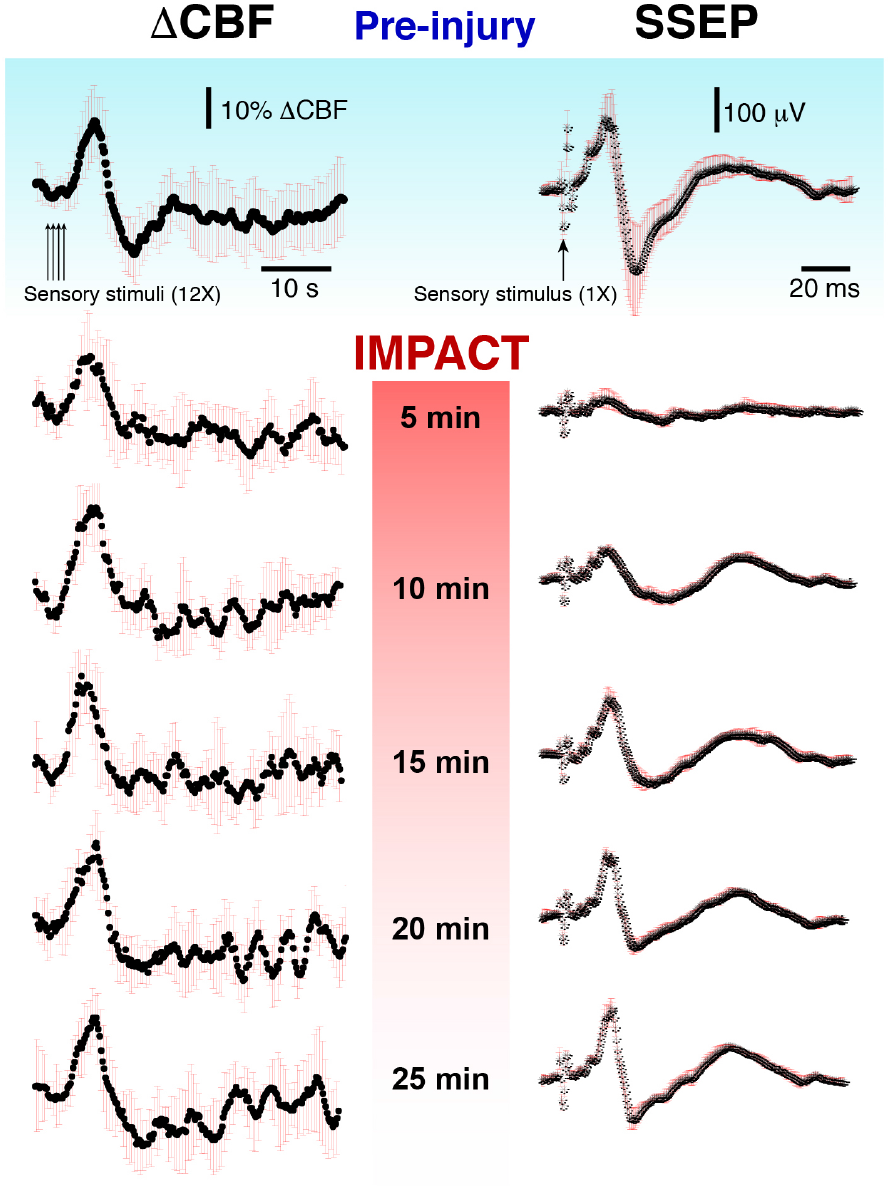
Sensory-evoked blood flow responses from forepaw stimulation for a representative experiment pre-and postinjury. The ΔCBF and SSEP traces shown here represent the average of multiple stimulations (average of 10 trials preinjury and 5 trials for each time point post-injury). The black line is the average and the red error bars represents a standard deviation.

**Fig. 4.**
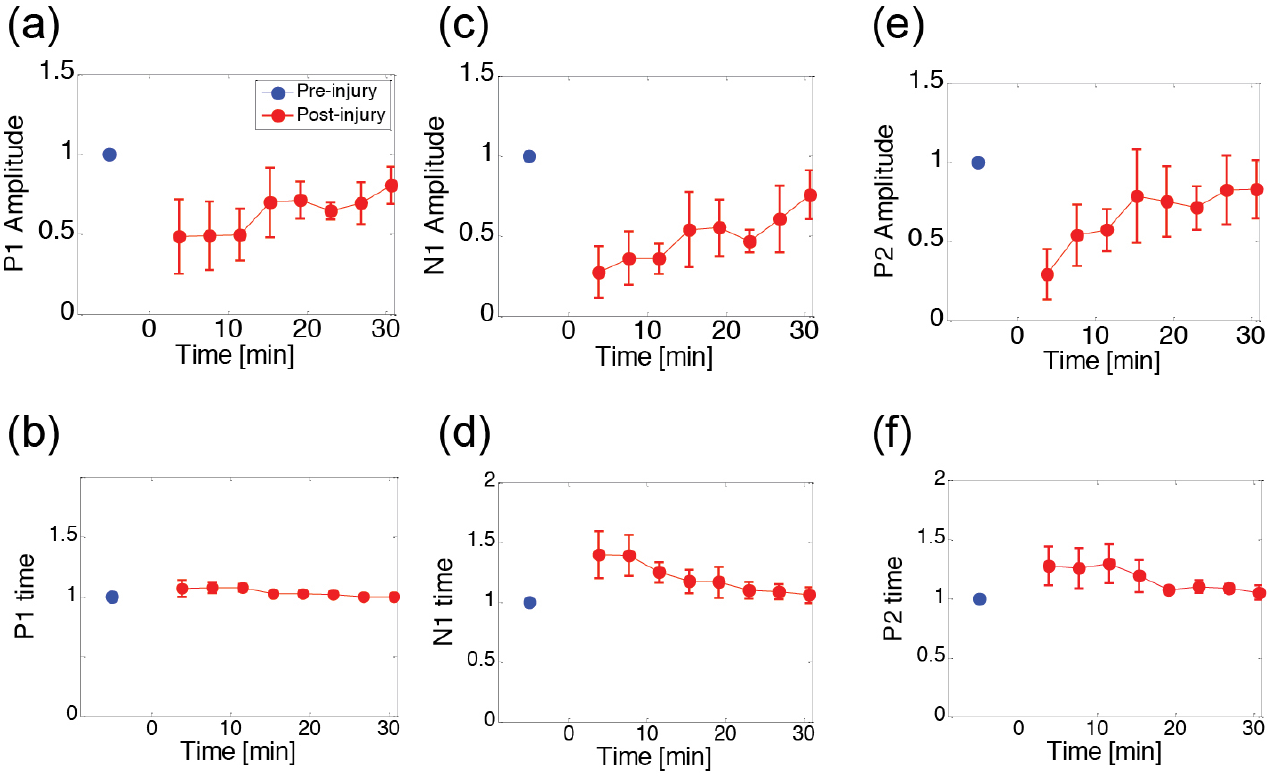
Grand average summary of the effects of CCI on the amplitude and latency of prominent time-domain features of the ΔCBF waveform. The red dots represent the average values over all animals (*n* = 6). Values are normalized to the baseline value (blue dot). The time of impact is designated here as *t* = 0 s. Error bars represent standard error.

SSEPs displayed similarly profound alterations within the first 5 min after CCI. As depicted in Fig. 5, P1 fell to ~50% of its pre-injury amplitude, however N1 and P2 exhibited greater reductions of 73 and 71%, respectively. CCI additionally altered the peak latencies. Although the

**Fig. 5.**
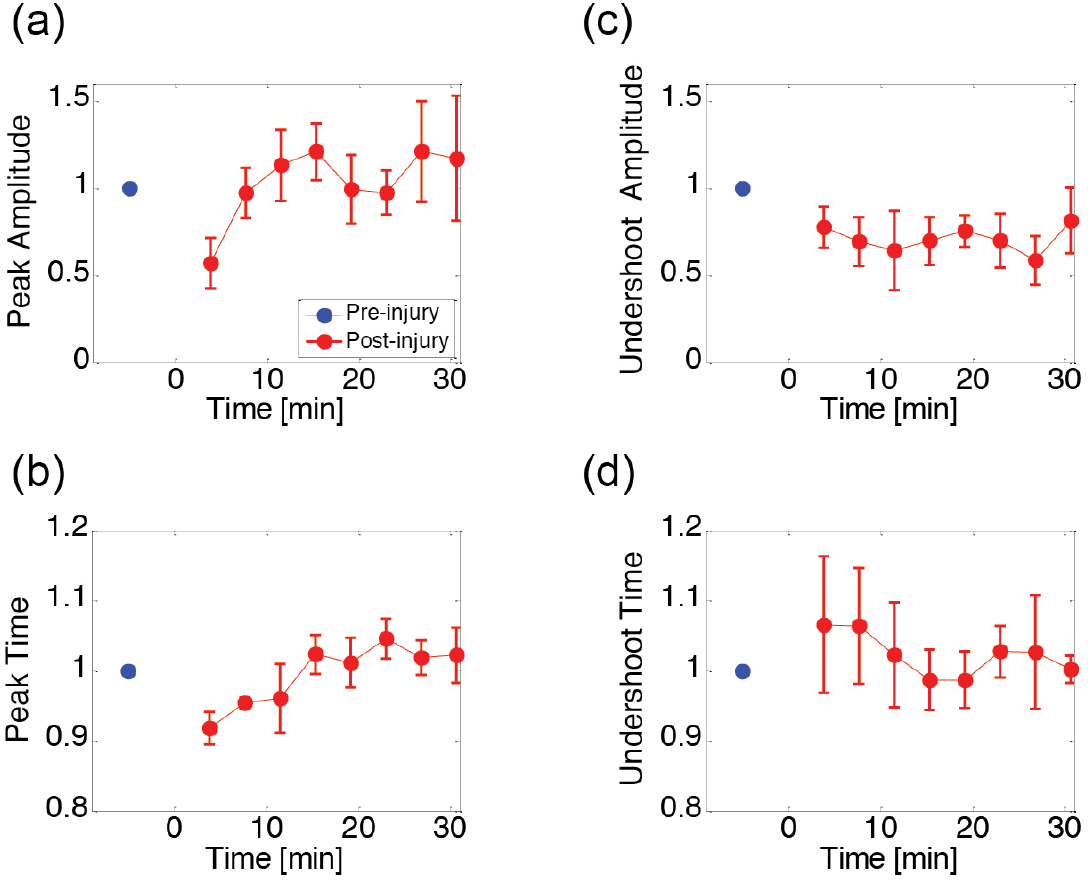
Grand average summary of the effects of CCI on major SSEP peaks. As in Fig. 4, red data points indicate post-injury averages over all animals (*n =* 6) ± S.E. and are normalized to baseline values (blue data points). The time of impact is designated here as *t* = 0 s.

P1 latency increased by less than ~7%, N1 and P2 both increased by upwards of ~40% and ~28%. In general, P1 appeared to be the least sensitive to injury, in terms of both amplitude and latency. In the ensuing 30 minutes, the peak latencies and amplitudes largely—although not entirely— returned to their baseline values, yet at differing rates. The amplitude of P2, for example, recovered more rapidly than N1 or P1 in the first 15 min. Overall, however, following CCI, the initial decrease in SSEP amplitude (peak-to-peak, N1 to P2) was significantly more profound than the drop in CBF. For example, SSEP peak-to-peak amplitude was reduced by nearly 75%, while CBF peak-to-peak dropped by only ~30%.

In addition to alterations in average amplitude and latency of SSEP and HRF features, injury acutely increased their trial-to-trial variability of these parameters (Fig. 6). In our time-domain measurements, we quantified this variability in terms of the coefficient of variation, *C_v_*, which is defined as the ratio of the standard deviation to the mean value. In the first five minutes following CCI, *C_v_* increased by over 100% for all electrophysiological and hemodynamic parameters except for the HRF undershoot amplitude, which increased by ~80%. Additionally, while the variability of other parameters returned to near baseline values within 10 minutes after injury, the HRF undershoot amplitude maintained an elevated variability over 30 minutes after injury, on average.

**Fig. 6.**
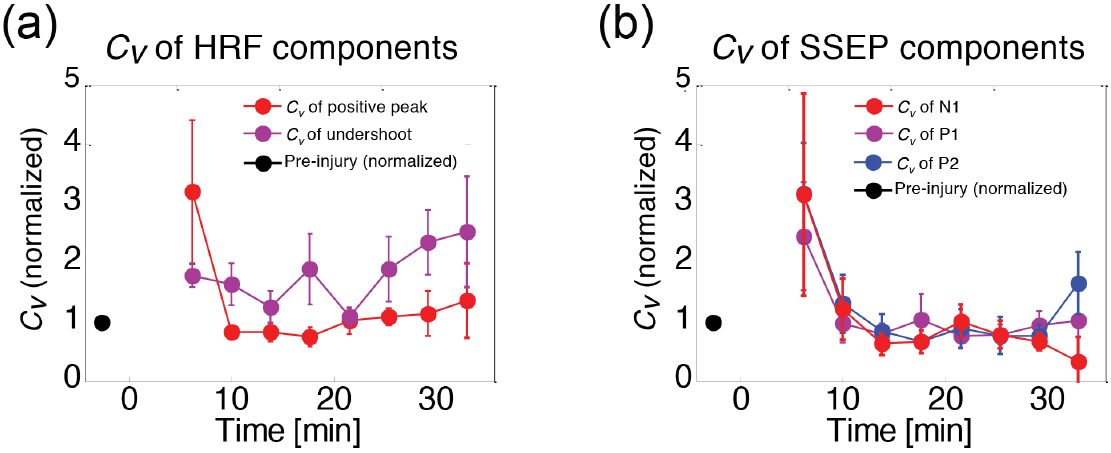
Alterations in the variability of evoked responses following injury. (a) Dynamics of the HRF variability are plotted over time, normalized to a baseline value (black dot). Injury was induced at 0 min in these plots. Variability is quantified here in terms of the coefficient of variation, *C_v_*, defined as the ratio of the mean value to the standard deviation. *C_v_* is plotted for both the initial HRF peak (in red) and the subsequent undershoot (purple). (b) Evolution of *C_v_* for SSEP peaks N1 (red), P1 (purple), and P2 (blue), normalized to baseline values (black dot).

### Spectrotemporal Analysis the Hemodynamic Response Function Following TBI

Modulation of prominent aspects of the hemodynamic response such as peak amplitude and latency are straightforward to quantify because the waveforms can be fit well to a conventional hemodynamic response function model (as described in MATERIALS AND METHODS). However, injury also induced new temporal features into the HRF. For example, in the ~30 s following each evoked peak in CBF there emerged slow oscillations in the HRF at a frequency of roughly 0.1 – 0.3 Hz. These features introduce additional peaks that cannot be effectively fit using the standard, two-gamma function model. As an alternative approach for quantifying these new components of the HRF, we performed time-frequency spectral analysis to explore and more directly visualize these effects. As depicted in Fig. 7, injury evoked changes in spectral power throughout the duration of the major peaks (increase and undershoot) of the HRF, as well as in the interim between stimulus sets. Although the apparent spectrotemporal changes development of temporally-separated “notches” in the spectral power for frequencies below 1Hz, we explored whether these changes were significant across animals by assessing the patterns of spectrotemporal differences from pre-injury values (Fig. 7b). The regions of statistically significant spectrotemporal changes in power are relatively sparse, however when summed over frequency and for all experiments, significant changes coincided with individual peaks in the “oscillatory” region, indicating a degree of phase-locking (Fig. 7c).

**Fig. 7.**
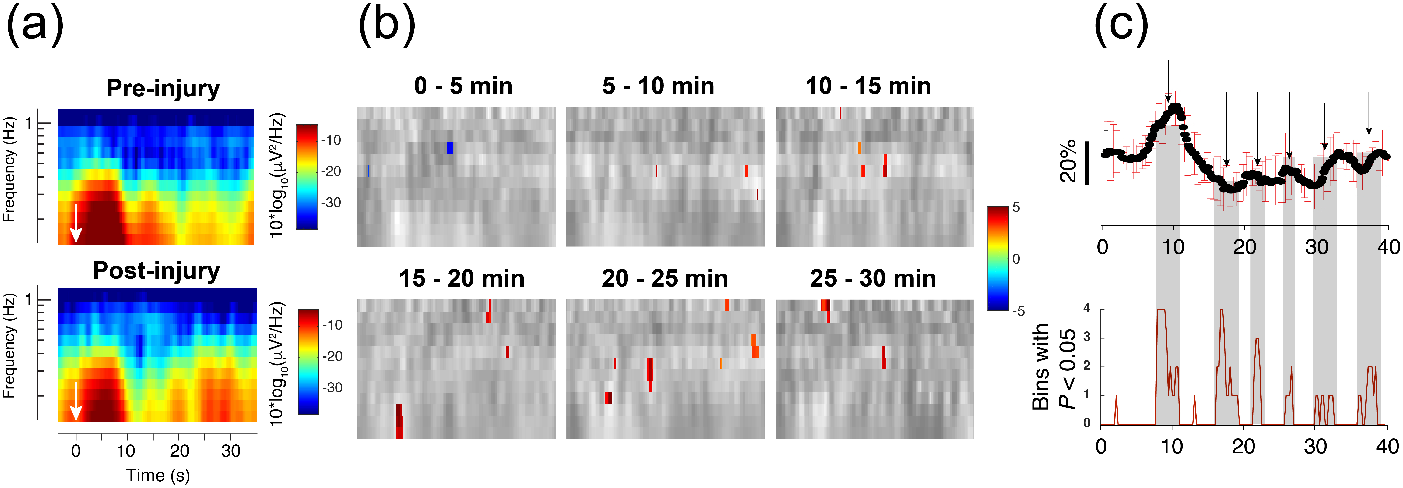
Time-frequency analysis reveals significant changes in spectrogram after injury. (a) Representative time-frequency spectrograms from a single animal showing the alterations in spectrotemporal content following CCI. The white arrows indicate the time point at which forelimb stimulation was applied. (b) Spectrotemporal changes over time, averaged over all animals. Each image depicts, in grayscale, the difference in spectrotemporal content via subtraction of the pre-injury spectrogram (darker shades indicate decreases, brighter shades indicate increases). Superimposed in color on the grayscale difference spectrogram are points that, when analyzed over all experiments, demonstrated a difference from the pre-injury values with P < 0.05 using a two-tailed Student’s t-test. The colors are defined as in the colorbar of (a), and note that nearly all of the spectrotemporal power changes are focal increases. (c) Comparison of a representative post-injury HRF (here, at a latency of 25 min following injury) with a plot showing the sum, over all experiments, of pixels showing alterations from baseline with P < 0.05. Note that the significantly changed spectrotemporal features lined up with the cycles of apparent “oscillations” following injury. The gray shading projected onto the post-injury HRF (a representative plot) highlights temporal periods that contained statistically-significant alterations in spectral power, assessed over all experiments (n = 6).

To quantify this apparent coherence effect, we analyzed the inter-trial coherence (ITC) of the spectrogram (Fig. 8), which reflects spectrotemporal phase-locking and variability among evoked responses [27]. Similar to the results of spectral power analysis depicted in Fig. 7, injury qualitatively changed ITC most prominently for regions following the initial HRF peak. A large fraction of the alterations in ITC were increases, most prominently in the period after the CBF undershoot and for frequencies above 0.4 Hz. The one region that displayed a significant decrease in ITC was that corresponding to the spectrotemporal representation of the CBF undershoot, indicating greater variability of that portion of the HRF after injury. Although significant spectrotemporal increases in ITC (P < 0.05 via two-tailed Student’s t-test) were apparent throughout the timecourse of the HRF, we investigated the temporal distribution of ITC alterations (Fig. 8c) by analyzing the HRF as a series of three “regions,” defined by physiologically relevant landmarks. When summed over frequencies, the period following the HRF undershoot (region three) demonstrated significant alterations from baseline.

### Correlations in the Dynamics of Cerebral Blood Flow and Electrophysiological Responses Following TBI

Given the diversity of signals that were sensitive to injury, we sought to merge the findings by exploring correlations among their recovery kinetics. The matrix in Fig. 9 presents correlation coefficients comparing the post-CCI quantitative dynamics depicted in Figs. 4 and 5. among the varied injury-sensitive metrics. Overall, SSEP peak amplitudes and latencies were highly intercorrelated. In comparison, the recovery kinetics of the HRF components were relatively weakly intercorrelated. Among the cross-correlation terms indicative of coupling between hemodynamic and neural activity, we identified two major trends which are highlighted in Fig. 9. Notably, of all time-domain electrophysiological features that we explored, the amplitude of the HRF peak was most significantly correlated with the amplitude of P2 (*z*-score > 3, based on the standard deviation of the distribution of Pearson’s correlation coefficients). Additionally, while P2 represented the strongest cross-modal correlation in terms of amplitude, the *latency* of the initial HRF peak was highly correlated with all aspects of the SSEP peaks.

**Fig. 8.**
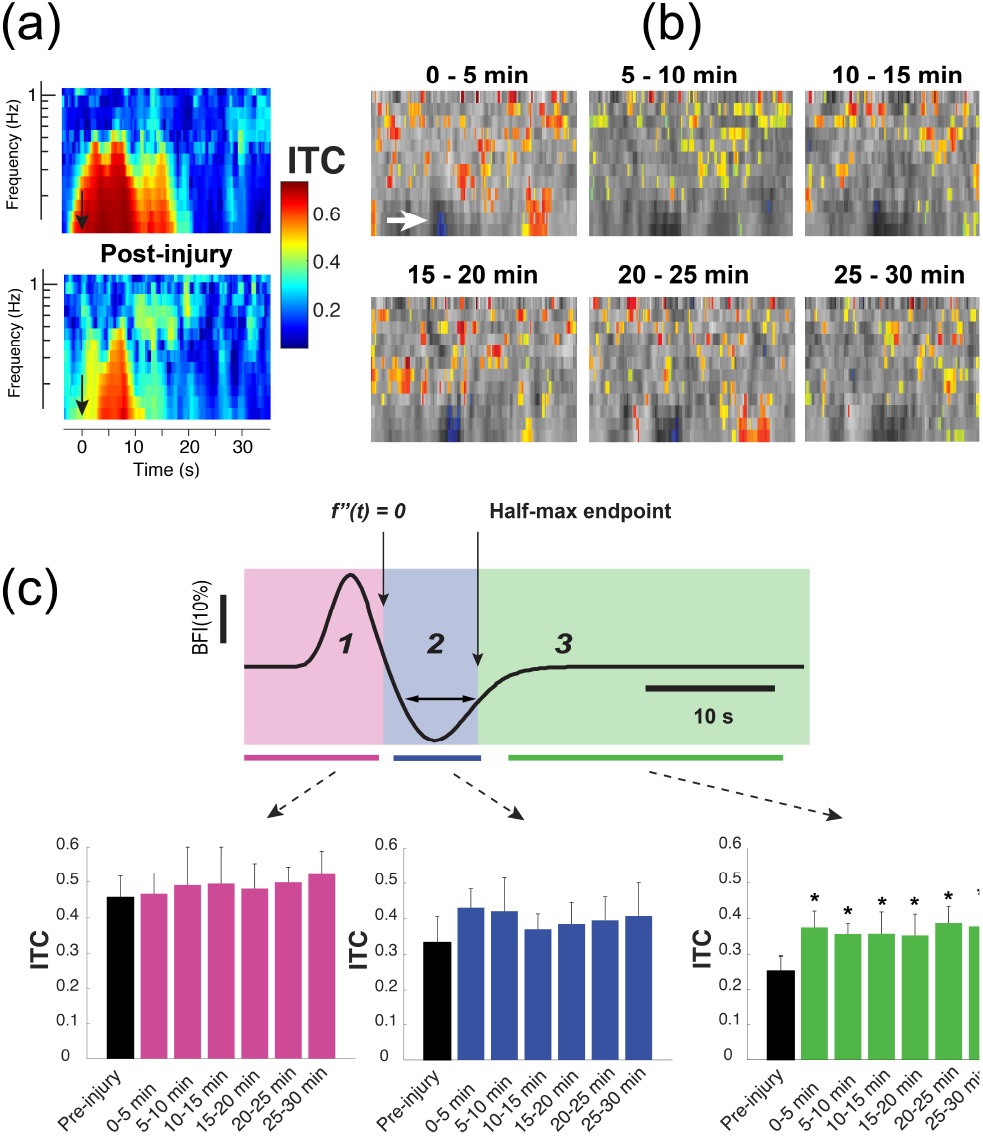
Inter-trial coherence alterations following controlled cortical injury. (a) Representative spectrograms depicting the absolute magnitude of inter-trial coherence (ITC) before (top) and after (bottom) injury. These particular spectrograms are derived from the same animal as in Fig. 7a. The color definitions are as indicated in the ITC colorbar; a value of 1 represents perfect, in-phase spectral responses. While the depicted values are real values, they represent the absolute magnitude of complex values that originally contain phase information. The black arrows indicate the time point at which forelimb stimulation was applied. (b) A series of ITC spectrograms, averaged over all animals, showing features that differ significantly from pre-injury with P < 0.05 obtained from a two-tailed Student’s t-test. As in Fig. 7b, the statistically significant pixels, which are color-coded according to the colorbar of (a), are superimposed on a grayscale spectrogram representing the post-injury ITC spectrogram from which pre-injury spectrogram values have been subtracted. While most of the changes represent increases in ITC at time points following the initial HRF peak and undershoot, there was a significant decrease in ITC in the region of the undershoot itself, roughly the area of frequencies below 0.4 Hz between 12 and 16 seconds post-stimulus (indicated by the white arrow in the top left superimposed image). (c) Regional assessment of ITC changes following injury. A sample HRF fit divided into three regions for assessing changes in ITC at particular phases of the functional hemodynamic response is shown on top. The specific regions were defined as (1), an initial “rise” extending from the time directly after sensory stimulation cessation up through the point where the second derivative of the HRF fit passes zero; (2) the undershoot period, extending from the end of (1) through the latest point of intersection of the HRF fit with the undershoot half-maximum (or minimum, here) line; (3) the period of time following the main two phases of the HRF and extending until the subsequent stimulus period. Asterisks in the right-most bar chart on the bottom indicate values that differ from baseline, pre-injury values with P < 0.05 according to a two-tailed Student’s t-test.

**Fig. 9.**
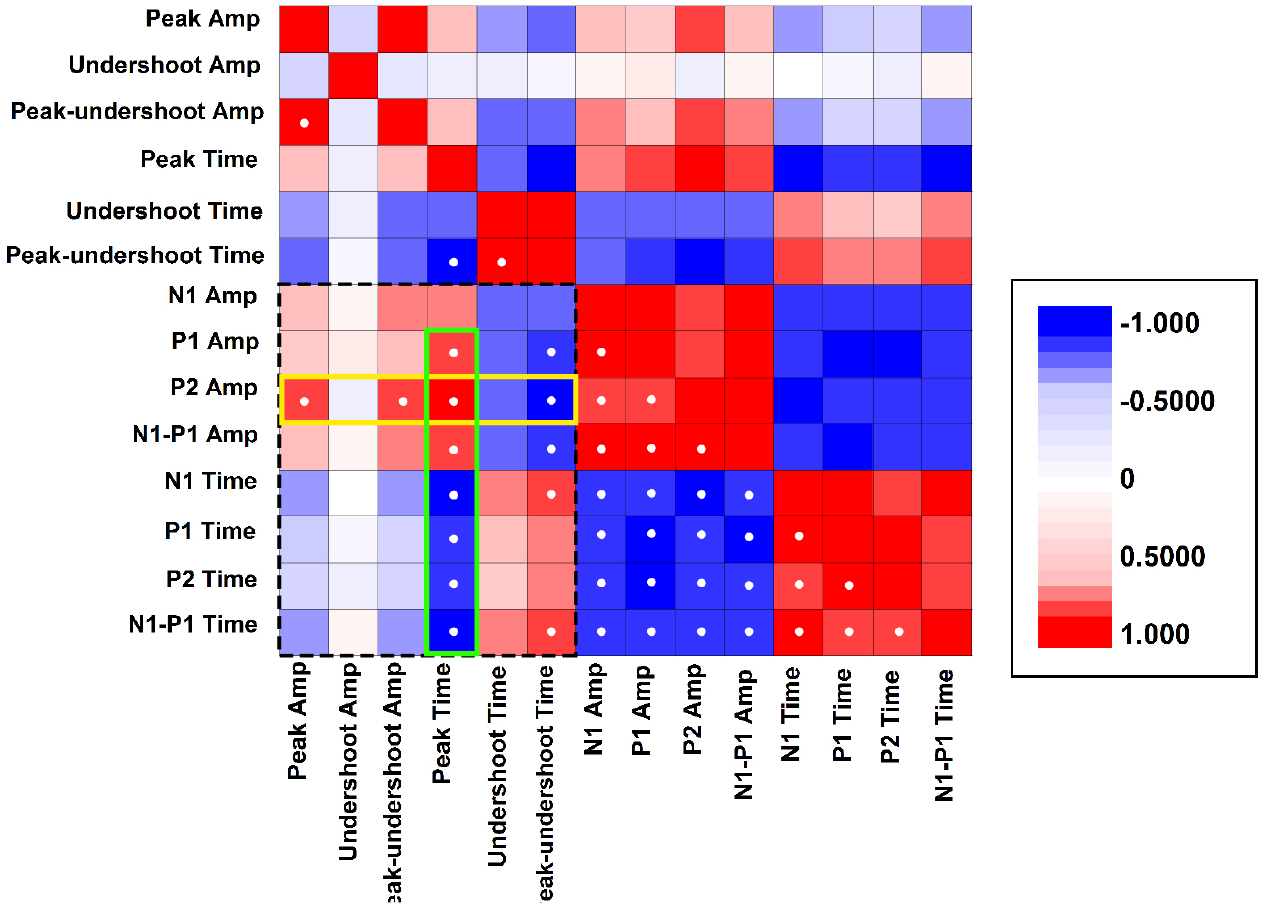
Correlation matrix comparing the post-CCI recovery trends for the prominent features of the CBF and SSEP responses. Each matrix element represents the Pearson’s correlation coefficient obtained by comparing the post-CCI time-course (as in Figs. 4 and 5) of the major peaks and latencies. A correlation coefficient value of 1 represents identical waveforms, -1 indicates that the two traces are perfectly anti-correlated. The matrix is symmetrical about the diagonal. Correlation values higher than 0.8 are indicated with white dots. The dotted rectangular perimeter delimits the matrix region representing correlation between the two modalities, optical and electrophysiological. The yellow perimeter highlights the high correlation between the P2 amplitude and post-injury hemodynamics, whereas the green perimeter highlights the fact that the HRF peak latency is highly correlated with all aspects of SSEPs.

## Discussion

A major repercussion of traumatic injury in the brain is an ensuing mismatch between oxygen supply and neural metabolic demand [28], [29]. The HRF and local tissue oxygenation ultimately reflect an interplay between CBF and changes in the cerebral metabolic rate of oxygen (CMRO2), both of which are susceptible to alteration following acute injury. The primary mechanical injury, for example, can alter the vascular network that delivers oxygen to the parenchyma and, simultaneously, can alter neural activity and metabolic rate through a wide spectrum of mechanisms, including mechanical shearing [30] or excitotoxicity due to damaged glia [31]. Our concomitant measurements of CBF and electrical activity provide a picture of the repercussions of injury on both “supply and demand” aspects of the networks underlying sensation.

While the general trends in SSEP alterations following injury are consistent with previous work in animal models [32]–[35], in the present study, we were able to identify time-domain alterations in the SSEP in finer detail. The most prominent SSEP peaks—N1, P1, and N2— exhibited differing alterations after injury, some of which are consistent with the known anatomical correlates of time-domain waveform. For instance, considering that the site of primary injury in CCI is the cortical gray matter [36], it is not entirely surprising that P1 (~13-ms onset delay pre-injury) exhibited the mildest decrease following injury given the fact that this component of the SSEP reflects subcortical activity related to thalamocortical relay [37]. Likewise, the high sensitivity of P2 likely reflects the fact that its underlying generators are derived from cortico-cortical processing, which is sensitive to lateral cortical perturbations. For example, modulating cortical activity with topical application of pharmacological agents generally affects N1, but largely spares P1 [38]. Consistent with this, in our previous work using a high-intensity focused ultrasound model for explosive blast, the mechanical insult penetrated deeper beneath the surface of the cortex and P1 was more significantly altered [35]. The high correlation between hemodynamics and P2 may also reflect an intrinsic, disproportionately large influence of superficial cortical injury on electrical activity and hemodynamics within the cortical microvasculature. The superficial layers of the cortex feature extensive lateral networking with other cortical areas [39], thus local disruption due to impact injury would likely have diffuse and far-reaching repercussions.

Within the temporal span of individual hemodynamic responses, the major features of the preinjury HRF that we observed match results obtained from PET [40], [41], arterial spin labeling (ASL) fMRI [42], [43], as well as optical investigations of flow [15], [44], [45]. The initial peak in the HRF reflects the influx of blood flow recruited by local changes in neural activity [46], and, on the timescale of days, its amplitude has been found to be reduced following traumatic injury [43] and stroke [47]. To our knowledge, however the impact of injury on sensory-evoked CBF, both in terms of amplitude and temporal waveform, has not been explored within the first moments following TBI. It remains to be seen whether our observed acute dynamics, which depict significant recovery of some aspects of the HRF, continuously evolve into the previously reported long-term deficits. In terms of injury-induced alterations in HRF peak timing, the same prior investigations reported a delayed HRF peak onset that is observable on the timescale of days following injury. Our observation of a reduced onset latency is novel, however it may be unique to the immediate aftermath of acute injury. In general, the fact that most aspects of the HRF and SSEPs substantially recover within a relatively short period of time is remarkable given the known chronic effects that TBI exerts on baseline hemodynamics, notably, changes in cerebrovascular autoregulation [48], [49]. The findings concur with previous reports that functional CBF is independent of baseline values [44], [50], [51], however our results indicate that this independence extends to injury-induced global hemodynamic changes.

It has long been assumed that the HRF undershoot is primarily a vascular phenomenon, representing a post-stimulus period of increased cerebral blood volume even after CBF and CMRO2 have returned to baseline [52]. However more recent work strongly indicates that the post-stimulus undershoot is highly dependent on neural activity [53]. Our results depict the postinjury undershoot as chronically attenuated and more variable. Similarly, SSEP components also displayed a marked increase in variability in both time and frequency domain. These observations are therefore consistent with the hypothesis that neural activity plays a significant role in shaping the undershoot. However, the fact that the variability and amplitude of the undershoot does not recover as fully as other HRF components even after SSEP attributes have returned to baseline status suggests that the prolonged undershoot alterations may not be primarily shaped by neural activity. Future studies involving systematic variation of stimulus parameters may be able to more directly probe this hypothesis.

Although we observed the emergence of prominent oscillations following injury (Figs. 7-8), such “ringing” in the functional CBF response has been reported in uninjured animals [54] and healthy humans [55]. However, there is no consensus on the physiological mechanisms that underlie this phenomenon. Models that capture the effect derive an oscillatory component based on inclusion of an inertial term, owing to blood volume in larger venules and arterioles [56]. Oscillations may also emerge from coupled molecular signaling pathways involved in the neurovascular coupling [57] or from cerebral vasomotion [58]. In either case, the effect involves a degree of coupling between local neural activity flow in more distal vessels. Given both the physical separation as well as differing mechanical properties of smaller vs. larger vessels, it is unlikely that injury exerts equal impact on both proximal and distal components of the neurovascular unit. Accordingly, a simplistic interpretation of the results is that the emergence of post-undershoot oscillations in the HRF reflects uneven perturbation of an intrinsically oscillatory system, regardless of whether the affected components are mechanical or phenomenological (e.g. related to molecular signaling). In terms of the physical paths of blood flow, the enhanced oscillatory signal may indicate “reflections” caused by an induced impedance mismatch between proximal and distal components of neurovascular communication.

A general challenge for parsing the bulk signals obtained through diffuse optical neuromonitoring techniques is that the spatial resolution is limited because the detected signals include light that has traversed a large span of optical paths in tissue [59]. In the case of DCS, although the signal origin is putatively weighted toward microvasculature, the fidelity with which the bulk signal aligns with a micro-scale portrait of blood flow in the layers of the cerebral cortex remains to be quantified. Future work in animal models of TBI that supplements DCS measurements with high-resolution imaging of activity in microvasculature, such as optical coherence angiography [60], [61] or two-photon microscopy [62], may better inform interpretation.

## Conclusion

While significant progress has been made toward understanding the repercussions and mechanisms of mTBI at the cellular level using invasive, high-resolution techniques, biomarkers for rapidly assessing human subjects in active settings outside the clinic are constrained to signals that can be obtained with noninvasive technology. Such measurement modalities, however, are limited in their ability to spatially resolve signals and biomarkers observed in preclinical work. As we have demonstrated, functional neurovascular coupling is a property that can be measured with low-resolution techniques and contains information about micro-scale physiological function. This information can be extracted from variable, high-noise physiological recordings because they are temporally synchronized with external stimuli that can be delivered in a controlled manner. The event-related optical and electrophysiological changes that we have discovered, albeit correlative, are significantly indicative of TBI in an animal model. We expect that these functional indicators will ultimately add informative dimensions to current, multimodal approaches for diagnosing mTBI in human subjects, given the heterogeneity of baseline values across populations.

## Acknowledgements

The mention of commercial products, their sources, or their use in connection with material reported herein is not to be construed as either an actual or implied endorsement of such products by the Department of Health and Human Services. The authors thank Drs. A. Parthasarathy, W. Baker, T. Durduran, A. G. Yodh, G. Yu, D. Busch, A. Koller, D. Ress, and J. Pfefer for helpful discussions, and Dr. S. Vasudevan for technical assistance. The work was supported by the National Science Foundation (Award 1641133) as well as intramural recruitment funds from New York Medical College.

